# Coronaviruses Spike glycoprotein endodomains: the sequence and structure-based comprehensive study

**DOI:** 10.1101/2023.08.16.553512

**Authors:** Prateek Kumar, Aparna Bhardwaj, Bodhidipra Mukherjee, Richa Joshi, Rajanish Giri

## Abstract

The flexibility of proteins makes them available to interact with many biomolecules in the cell. Specifically, such interactions in viruses help them to perform more functions despite having a smaller genome. Therefore, these flexible regions can be exciting and essential targets to be explored for their role in pathogenicity and therapeutic developments as they achieve essential interactions. In the continuation with our previous study on disordered analysis of SARS-CoV-2 spike cytoplasmic tail (CTR), or endodomain, here we have explored the disordered potential endodomains of six other coronaviruses using multiple bioinformatics approaches and molecular dynamics simulations. Based on the comprehensive analysis of its sequence and structural composition, we report the varying disorder propensity in endodomains of spike proteins of coronaviruses. The observations of this study may help to understand the importance of spike glycoprotein endodomain and creating therapeutic interventions against them.

## Introduction

Coronaviruses are enveloped, single-stranded, positive sense RNA viruses with genome sizes ranging from 27 to 32 kilobases, making them one of the largest RNA viruses ^1,2^. The coronaviruses belong to the *Coronaviridae* family and are classified into four subfamilies: alpha, beta, gamma, and delta ^3^. So far, the seven emerged human coronaviruses belong to alpha and beta coronavirus subfamilies. Among these, the alpha subfamily constitutes two human coronaviruses, namely 229E and NL63, while the beta subfamily constitutes the rest five, i.e., OC43, HKU1, SARS-CoV-1, MERS-CoV, and SARS-CoV-2 ^4^.The first reported strains (229E and OC43) of human coronaviruses were discovered in the mid-1960s and were reported to have caused only milder infections ^4^. Whereas five other strains have a pathogenic tendency towards humans, these include NL63, HKU1, SARS-CoV-1, MERS, and SARS-CoV-2 ^4^. Specifically, bats and rodents act as reservoirs of betacoronaviruses, but, in certain instances, they may get transferred to humans via spill over events ^5,6^. According to CDC, the total number of reported cases of SARS-CoV-1 was 8,096 during its outbreak. As per WHO, nearly 600 million cases of SARS-CoV-2 have been reported, and 2,585 cases of MERS have been reported as of 30^th^ August 2022^7^. 229E, OC43, NL63, and HKU1 are four other human coronaviruses commonly affecting people worldwide. However, information regarding infected people is not available for these HCoVs.

The coronaviruses show roughly spherical morphology with surface projections called spikes ^8,9^. This spike glycoprotein exhibits a trimeric complex before its attachment to the host cell membrane that aids in the fusion of viral membrane. It consists of two subunits S1 and S2, where S1 mediates the binding with the host cell membrane, whereas S2 mediates the fusion of viral and host membranes ^10^. Further, the S2 subunit consists of a fusion peptide (FP), two heptad repeat regions (HR1 and HR2 domain), a downstream transmembrane anchor domain, and a short cytoplasmic tail (CT) ^11,12^. Multiple cysteine residues flank the transmembrane segment of spike at its C-terminal (**Figure 1**). This domain lies in the intravirion region and contains multiple conserved cysteines suggesting its role in viral-host cell fusion. These cysteine residues have been reported to undergo post-translational modification like palmitoylation, inhibition of which can cause changes in infectivity and pathogenicity of the virus ^13,14^. However, few studies are related to this cytoplasmic tail or endodomain of spike protein.

**Figure 1:**
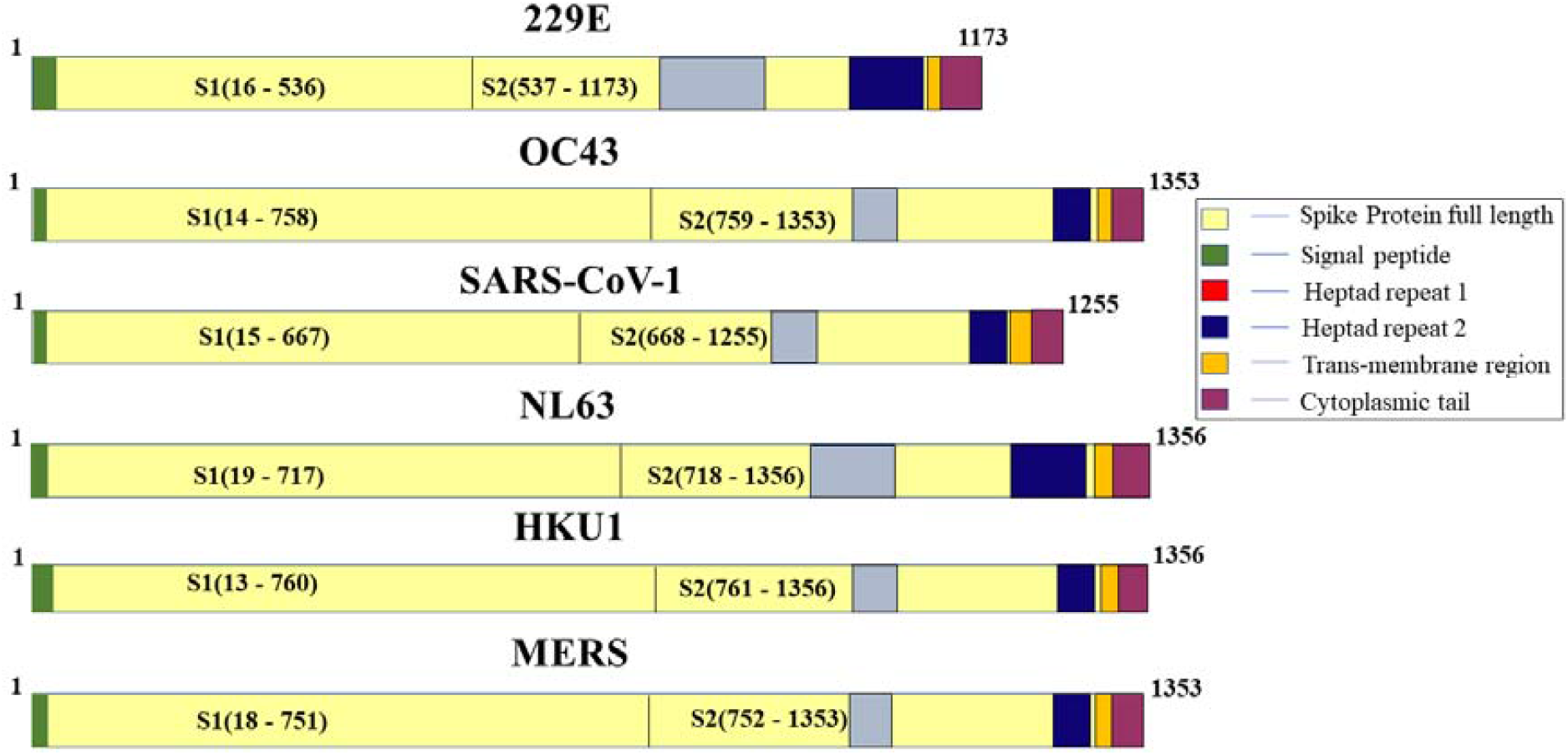
Schematic representation of literature-based characterized domains of spike glycoproteins of six coronaviruses.

Recently, we have identified that the SARS-CoV-2 spike CTR or endodomain is an intrinsically disordered protein region and may undergo a structural transition in the presence of other biomolecules under physiological conditions ^15^. Hence in this report, we are trying to answer the following question about the conformational ensembles of endodomain of spike protein of other six coronaviruses. Are they all disordered like SARS-CoV-2 spike endodomain? Or are there differences in the fold and dynamics of spike endodomains of different coronaviruses? Unfortunately, no single crystal, cryo-EM or NMR study is available that details the fold of endodomain of spike protein of all these coronaviruses. Therefore, we used various online predictors to analyze the spike cytoplasmic region’s disorder propensity. Further, we modelled the region in-silico via AlphaFold2. In isolation, we performed extensive MD simulations to get a deeper insight into the structure of the SARS-CoV-2 spike endodomain.

## Results

### Endodomains sequence retrieval and Multiple sequence alignment

Since the endodomains are extremely important for transporting spikes into Golgi apparatus via COP-I vesicles for post-translational modifications, endodomains can be promising targets through druggable inhibitors. Therefore, to begin with, the study of CTRs, all sequences were retrieved from full-length spike S2 subunit sequences from UniProt **(Table 1)**. Also, all the available structures in PDB did not possess transmembrane and endodomain in the structures **(Table 1)**. Therefore, based on the topology given on UniProt, the cytoplasmic regions were selected and used for further study.

**Table 1:** UniProt IDs and residues for the cytoplasmic regions of spike glycoproteins from all seven coronaviruses used in this study. The available spike structures of different domains associated with mentioned UniProt IDs are listed out in the table. The notable point is for all coronaviruses, the endodomain structures are not deciphered so far.

| HCoV | UniProt ID | Cytoplasmic Residues | Available spike structures (PDB IDs) |
| --- | --- | --- | --- |
| **229E** | P15423 | 1136-1173 | **7** (5YL9, 5ZHY, 5ZUV, 6ATK, 6U7H, 7CYC, 7CYD) |
| **OC43** | P36334 | 1319-1353 | **1** (7M51) |
| **SARS-CoV-1** | P59594 | 1217-1255 | **13** (1WNC, 1WYY, 1ZV7, 1ZV8, 1ZVA, 1ZVB, 2AJF, 2BEQ, 2BEZ, 2DD8, 2FXP, 2GHV, 2GHW) |
| **NL63** | Q6Q1S2 | 1317-1356 | **5** (2IEQ, 3KBH, 5SZS, 7FC3, 7KIP) |
| **HKU1** | Q5MQD0 | 1322-1356 | **2** (5GNB, 5KWB) |
| **MERS** | K9N5Q8 | 1318-1353 | **13** (4L72, 4MOD, 4XAK, 4ZPT, 4ZPV, 4ZPW, 5GMQ, 5GR7, 5GSB, 5GSR, 5GSV, 5GSX, 5VYH) |
| **SARS-CoV-2** | P0DTC2 | 1235-1273 | **13** (6LVN, 6LXT, 6LZG, 6MOJ, 6M17, 6M1V, 6VSB, 6VW1, 6VXX, 6VYB, 6W41, 6WPS, 6WPT) |

The literature on spike glycoprotein has revealed the presence of a cysteine-rich motif flanking the transmembrane domain ^1,13^. This abnormally high number of cysteine residues near the transmembrane domain is conserved in most coronavirus S proteins ^13^. As per the MSA results, coronavirus spike endodomains revealed the presence of a cysteine-rich motif with multiple conserved cysteine residues in all studied coronaviruses **(Figure 2)**. Another identifiable characteristic is the presence of an endoplasmic reticulum (ER) retrieval signal located at the end of each endodomain sequence. Following previous literature, it has a conserved dibasic character containing a lysine (K) and histidine (H) residue in orientation KXHXX or KXXHXX ^16^. It may help newly synthesized proteins in the aspect of subcellular localization. In OC43, 229E, and NL63, the YXXØ motif, lysosomal localization signal is also present as YQEL, YYDV, and YYEF, respectively ^17^.

**Figure 2:**
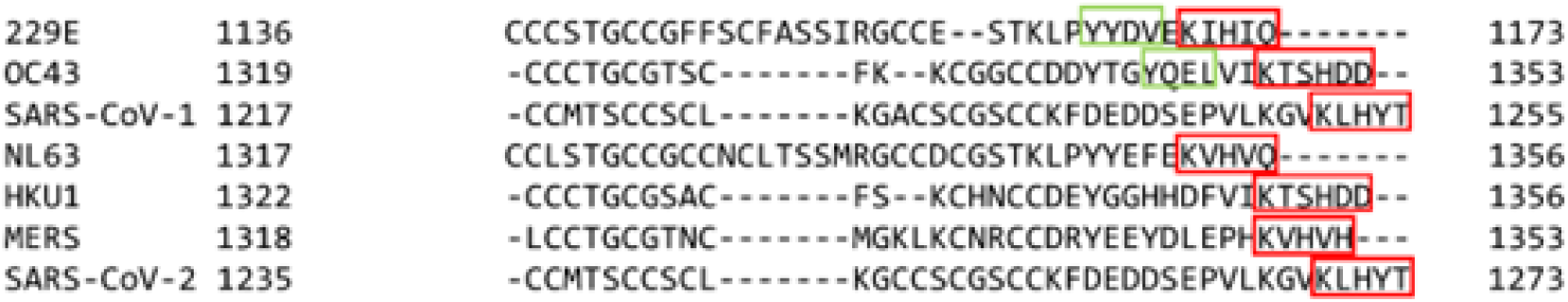
Multiple sequence alignment of Spike intravirion regions. With rectangle in red and green colour highlighting the dibasic motif, KXHXX or KXXHXX, and YXXØ motifs respectively.

### Intrinsic Disorder propensity

In the previous study, through computational and experimental approaches, we identified that the intravirion region or endodomain of SARS-CoV-2 is intrinsically disordered ^15^. This study led us to think about our next curiosity about the conformations of endodomains of the other six coronaviruses. Hence, we analyzed the disorder propensity of spike cytoplasmic domain of six other human coronaviruses. We used multiple disorder predictors, including the PONDR family, IUPred2A, PrDOS, and DisEMBL. For all six viruses, redox plus showed the least disorder propensity along with PrDOS and VLXT **(Supplementary Figure 1)**.

As observed from our results based on PPID, the disorder propensity of 229E has the least propensity, whereas SARS-CoV-1 is the highest for disorderness. (**Figure 3**). According to the mean value, most residues have high disorder propensity while some have moderate (1231-1242). Furthermore, in NL63, HKU1, OC43, and MERS, most residues showed moderate disorder propensity, while in 229E, most residues have low disorder propensity (residues 1140-1168). IUPred2A with the redox-state function was used as the cytoplasmic region is cysteine-rich. The redox minus state, where all cysteines are replaced by serine (as in reducing conditions, Cys behave as a polar amino acid similar to Ser), shows high disorder propensity for all cytoplasmic spike regions except for 229E, where redox minus state shows most of the residues to have low disorder propensity (residue 1143-1165) **(Supplementary Figure 1)**. In SARS-CoV-2, the overall disorder predisposition was observed to be nearly 90% for residues 1242-1273 which is highest among all seven coronaviruses ^15^.

**Figure 3:**
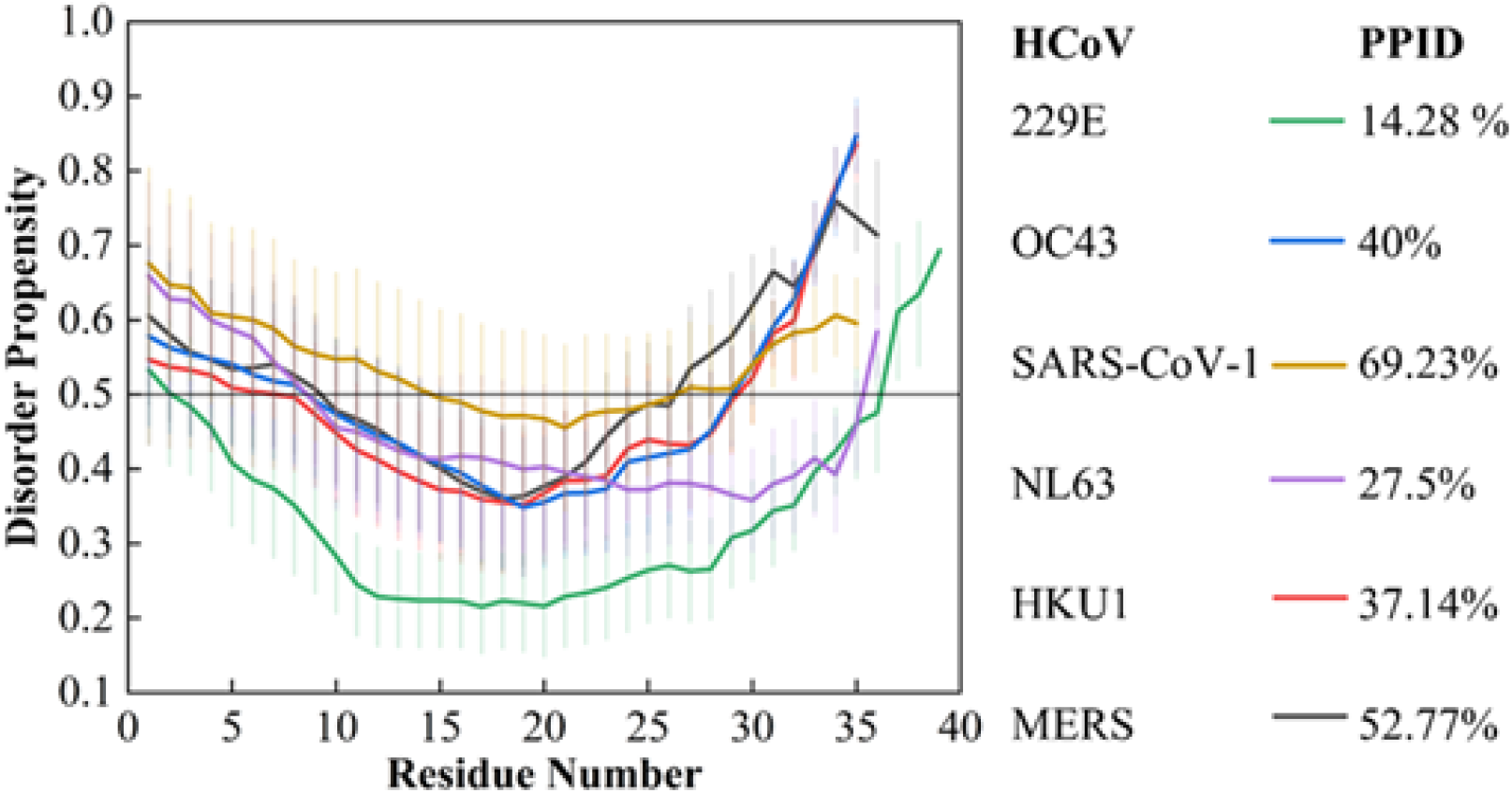
Disorder predisposition of spike cytoplasmic tails of human coronaviruses: Disorder propensity and percent of predicted intrinsic disorder (PPID) of spike cytoplasmic tails of 229E, OC43, SARS-CoV-1, NL63, HKU1, and MERS. Reference line on 0.5 score shows cut off for disorderness. The region above it is considered as disordered. The vertical bars on mean represent standard error of mean.

### Molecular Recognition Features (MoRFs) and SLiM (Short Linear Motifs) Predictions

Viruses possess short and long motifs that help them recognize and hijack mammalian host pathways. The disordered proteins of viruses contain these motifs, aiding their interaction with the host ^18^. Due to their tendency to change hosts, it is faster to evolve SLiMs compared to fully structured domains for carrying out viral-host interactions. This variability of final conformation aids in the adaptability of the protein by altering its surface area of interaction by presenting new motifs ^19–23^. Therefore, we have also investigated the motif recognition sites in all coronaviruses’ endodomain of spike proteins.

MoRFChibi predicted significant spike regions cytoplasmic regions of all six coronaviruses to be MoRFs, as tabulated in **Table 2**. However, according to the ANCHOR web server, no MoRF regions exist in any spike cytoplasmic regions. Furthermore, to get insight into regions that may function as motifs, we used the ELM database to identify short linear motifs in the spike endodomain sequences **(Figure 4)**. The complete functions and predicted SliMs for all the spike cytoplasmic regions are shown in **Supplementary Table 1**. SLiM data made some exciting revelations about spike endodomains of the six coronaviruses, as explained below:

**Table 2:**
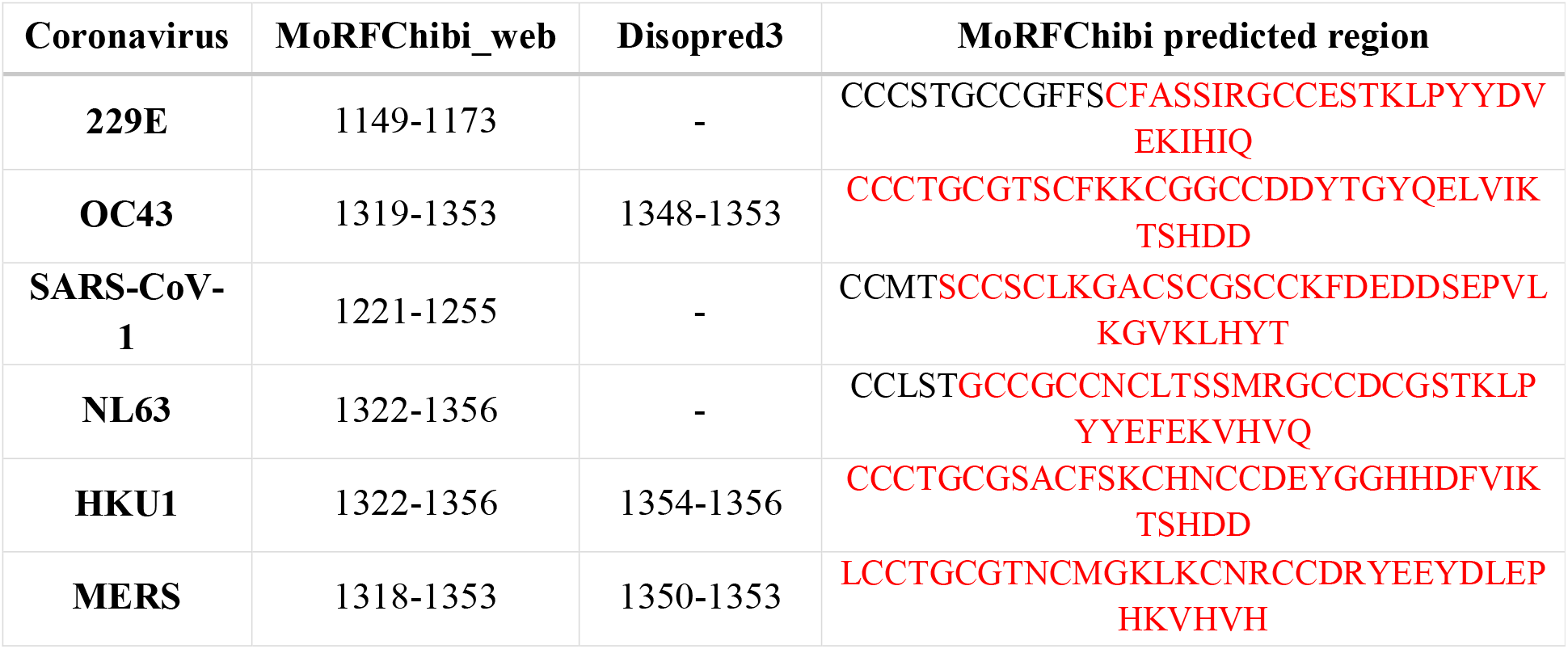
Prediction of MoRF regions through MoRFChibi and Disopred3. Red highlighted sequence represent MoRF regions of Human coronaviruses as predicted by MoRFChibi.

**Figure 4:**
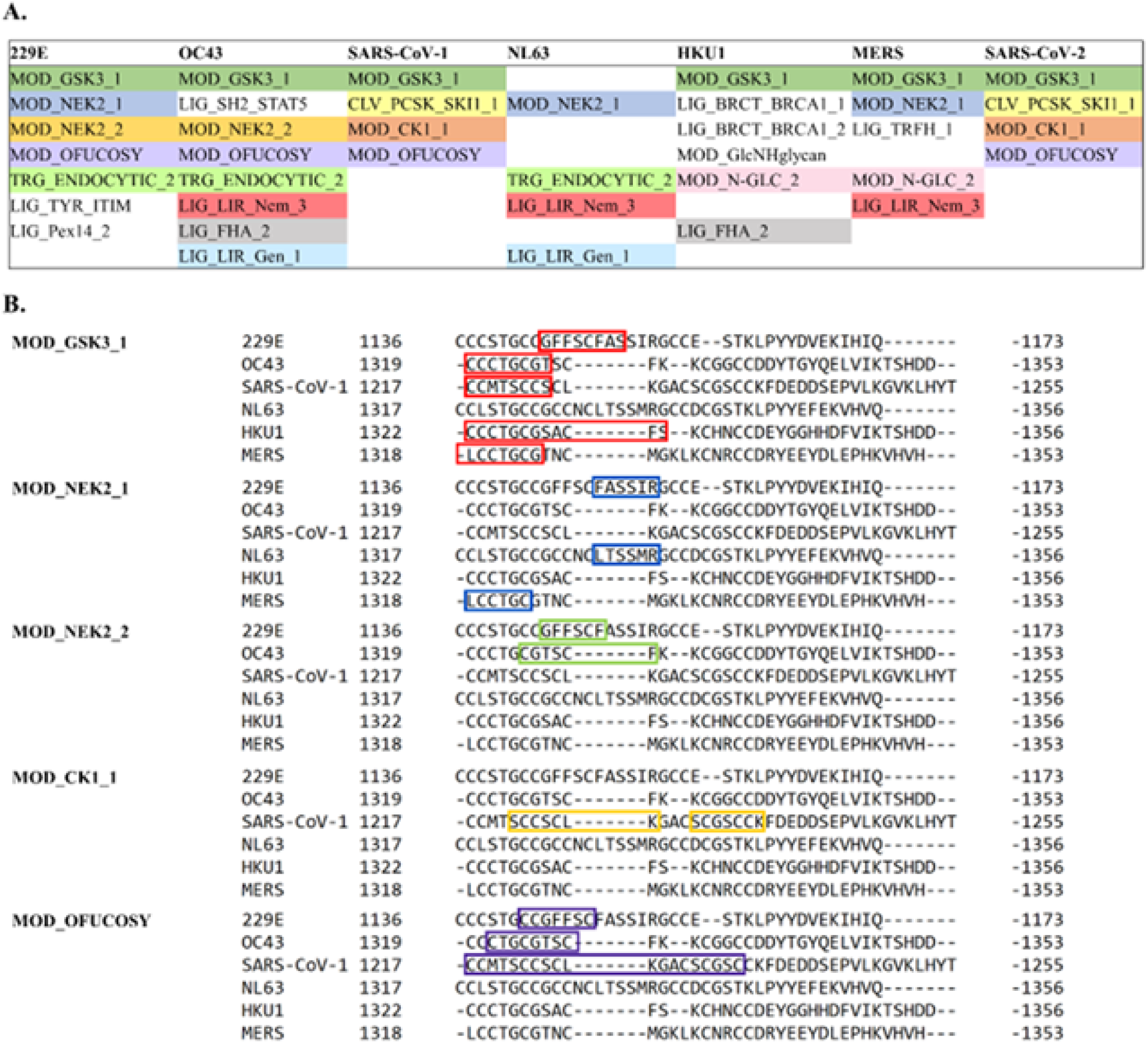
Predicted SliMs for spike cytoplasmic regions from eukaryotic linear motif (ELM) server: [A] ELM data represented according to timeline of occurrence of human coronaviruses. **[B]** Eukaryotic linear motif sequences as predicted by the ELM server (rectangles represent the identified motif sequences).

### Phosphorylation motifs

Protein phosphorylation refers to reversible post-transgression modification (PTM), in which a protein kinase attaches a phosphate group covalently to an amino-acid residue ^24^. The most commonly phosphorylated residues are Serine (S), Threonine (T), and Tyrosine (Y) ^25,26^. Phosphorylation may cause activation or deactivation of the known or unknown function of the protein and therefore plays a significant role in several biological processes like membrane transport, protein-protein interaction, protein degradation, enzyme regulation, cellular signaling ^27,28^, etc. According to SLiM information, the endodomain sequences tend to have various phosphorylation motifs near the transmembrane end of the C-terminal. These include GSK3 phosphorylation recognition site, MOD_GSK3_1; NEK2 phosphorylation sites MOD_NEK2_1, MOD_NEK2_2; and Casein kinase one phosphorylation sites MOD_CK1_1. It would be essential to note that these phosphorylation sites are present near the cysteine-rich end of the extreme C-terminal region.

According to the literature on SARS-CoV-1 and SARS-CoV-2, it is evident that cysteine residues near the transmembrane domain of CTR get palmitoylated ^13^. Palmitoylation of these residues is also vital for viral infectivity and other pathogenic characteristics like cell-to-cell fusion and cell surface expression of S protein ^13,29^. Palmitoylation involves the covalent attachment of palmitic acid to specific residues, typically cysteine, catalyzed by particular enzymes, increases the hydrophobicity of nearby regions, and therefore helps in subcellular localization, trafficking, and interaction with other proteins ^29^. Hence, palmitoylation might play a role in the trafficking of S protein to their designated sites. Previous studies of specific human proteins revealed reciprocal palmitoylation and phosphorylation of the proteins influencing transport ^30^.

### Fucosylation site on Spike endodomain

Protein glycosylation is a widespread post-translational modification responsible for controlling critical biological processes. Glycosylation of viral proteins has been extensively studied and revealed to play a crucial role in their infectivity and pathobiology ^31,32^. SARS spike proteins are no exception and have multiple glycosylation sites on the S protein. Most of these are in the form of oligo-mannose-type glycans ^33^. However, the function of this glycosylation in the pathobiology of coronaviruses is not entirely clear. Here in our study, ELM results revealed the presence of multiple fucosylation residues in the endodomains of most human coronaviruses, which is significantly more prevalent in both SARS species. Hence it can be assumed to play a role in their increased infectivity or pathogenicity.

Furthermore, to confirm the ELM predictions and identify the particular residues for PTMs, we utilized an AI-based predictor, MusiteDeep. Based on the outcomes, it was observed that the cysteine residues are susceptible to palmitoylation in every CoV **(Figure 5)**. Additionally, several other residues have been predicted to undergo modification post-translationally, such as lysine in 229E, OC43, SARS-CoV-1, NL63, and SARS-CoV-2 for acetylation (ac) as well as for SUMOylation in OC43, HKU1, and ubiquitination in SARS-CoV-1 & 2. Similarly, six types of modifications are identified in the endodomain of spike proteins of all CoVs.

**Figure 5:**
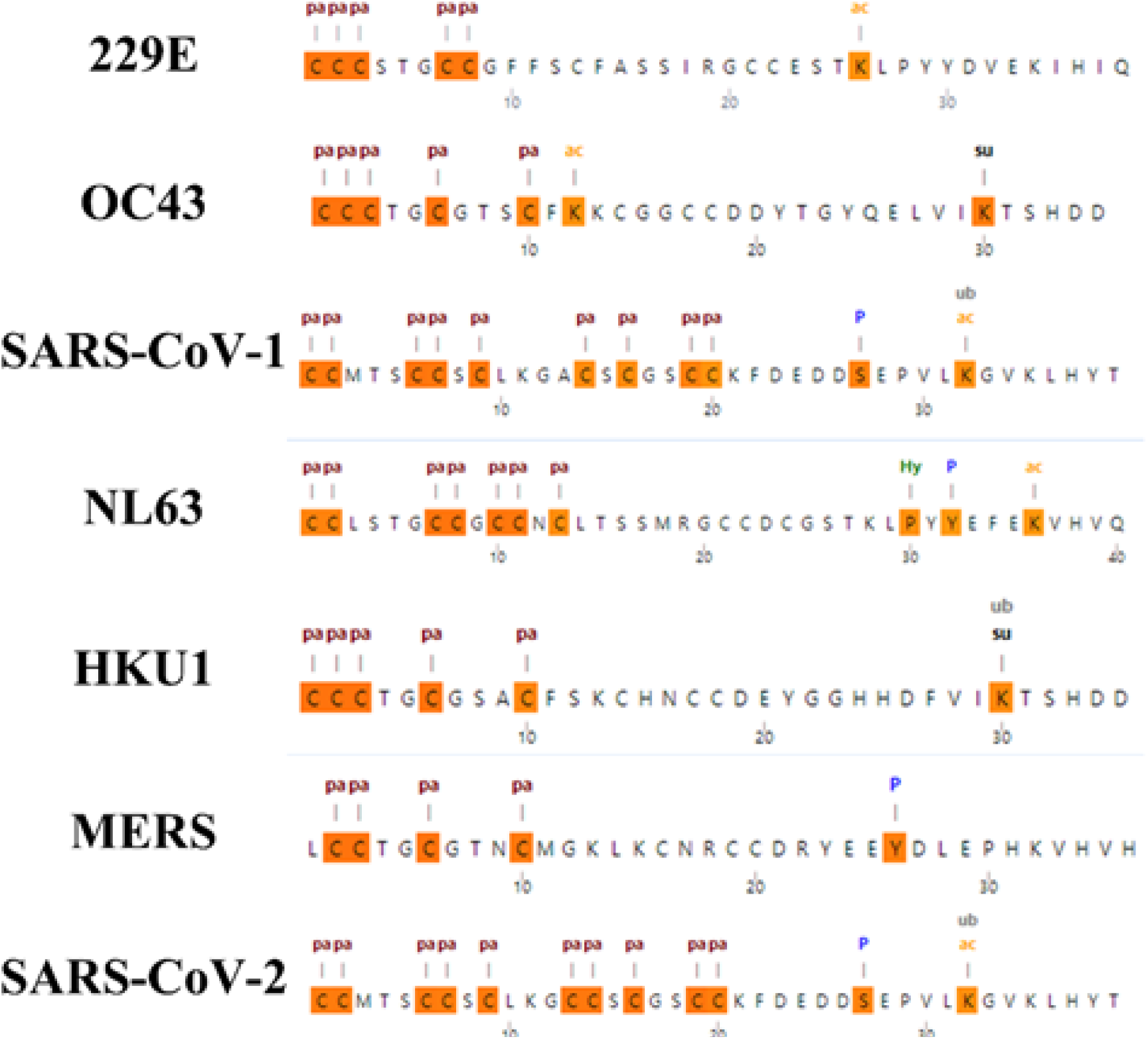
Snapshots of *MusiteDeep* server prediction results demonstrating the residues for post-translation modifications of all coronaviruses (pa-palmitoylation, ac-Acetylation, su-SUMOylation, P-Phosphorylation, ub-Ubiquitination, and Hy-Hydroxylation).

### Structure modelling

The experimentally determined 3D structures of spike glycoproteins have shown missing electron densities for the cytoplasmic regions. However, the modelled structures of full-length spike proteins have also shown similar outcomes. The endodomains in full-length spikes are modelled as mostly disordered structures having no or very less α-helical propensity **(Figure 6)**. Then, to decipher the fate of the endodomain in isolation, we modelled the spike protein endodomain of all six coronaviruses using AlphaFold2 **(Figure 7)**. According to the 3D structures formed by AlphaFold2, the residues 1338-1340 in MERS and 1225-1227 in SARS-CoV-1 form a small helix, whereas residues 1341-1345 and 1331-1335 in HKU1 form two small helices. Further, in NL63, 229E, and OC43, there are significant contributions of helices (10-13 residues) and disordered regions. In NL63, residues 1327-1337; in 229E, residues 1145-1159 (as per disorder prediction too for these residues, the mean disorder propensity is observed to be very low), and in OC43, residues 1325-1330 and 1332-1343 form helices (**Figure 7**). As calculated using the *2Struc* web server, all the spike cytoplasmic tails of CoVs, 229E, OC43, SARS-CoV-1, NL63, HKU1, and MERS had 44.7%, 45.7%, 7.7%, 27.5%, 31.4%, and 8.3% of total helix propensity, respectively **(Figure 8)**. As per the previous study, the spike endodomain of SARS-CoV-2 contained more than 80% of disordered regions. Here, the outcomes of modelled structures suggest SARS-CoV-1 has the highest percentage of the unstructured region, followed by MERS amongst all six coronaviruses. We have performed microsecond-level MD simulations with these modelled structures for all six studied CoVs.

**Figure 6:**
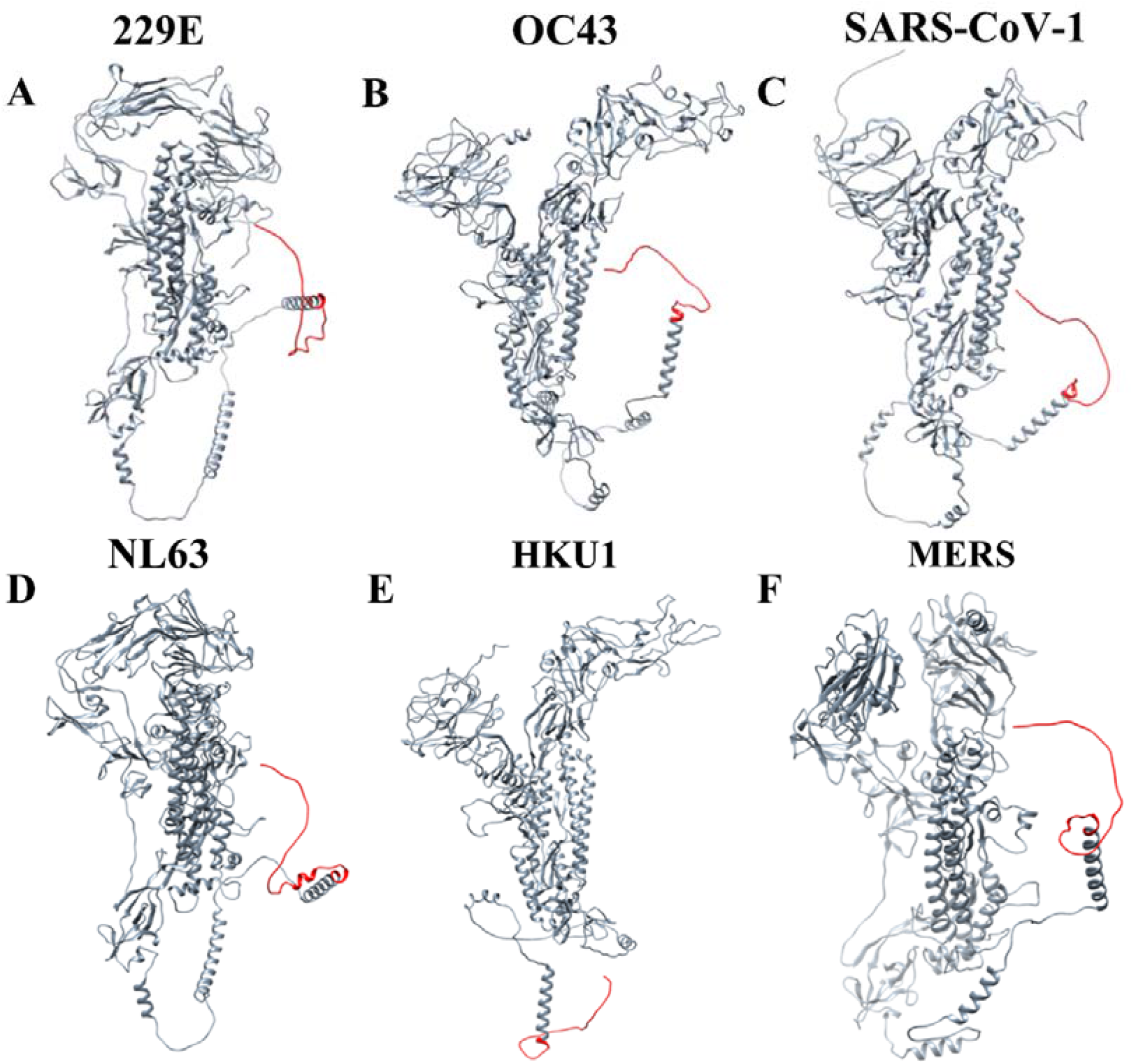
AlphaFold2 modelled 3D structures of spike full-length proteins. The endodomains are highlighted in red.

**Figure 7:**
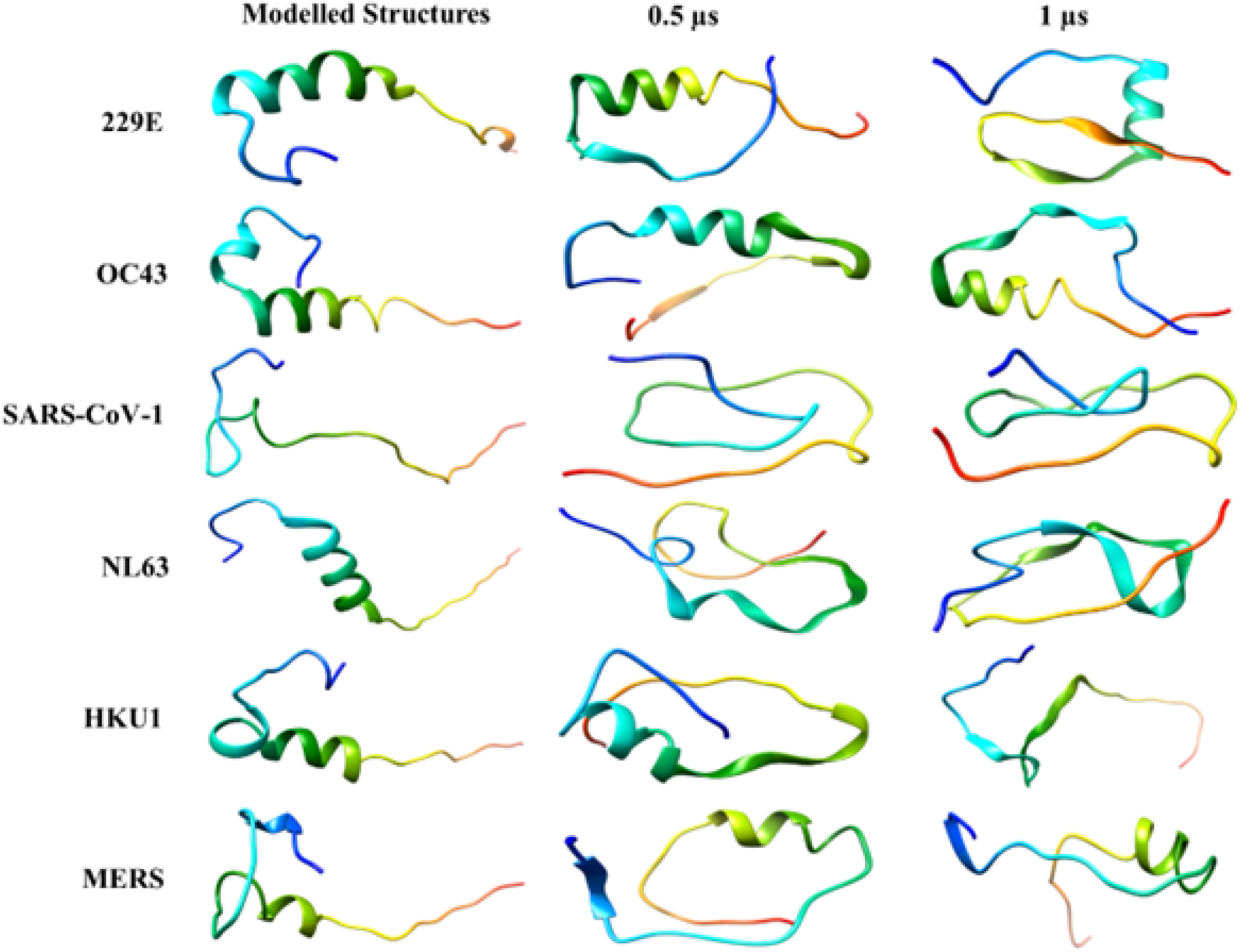
AlphaFold2 modelled 3D structures of spike endodomains in isolation along with 0.5 μs (second column) and 1 μs (third column) simulated structures of 229E, OC43, SARS-CoV-1, NL63, HKU1, MERS, respectively, from top to bottom rows.

**Figure 8:**
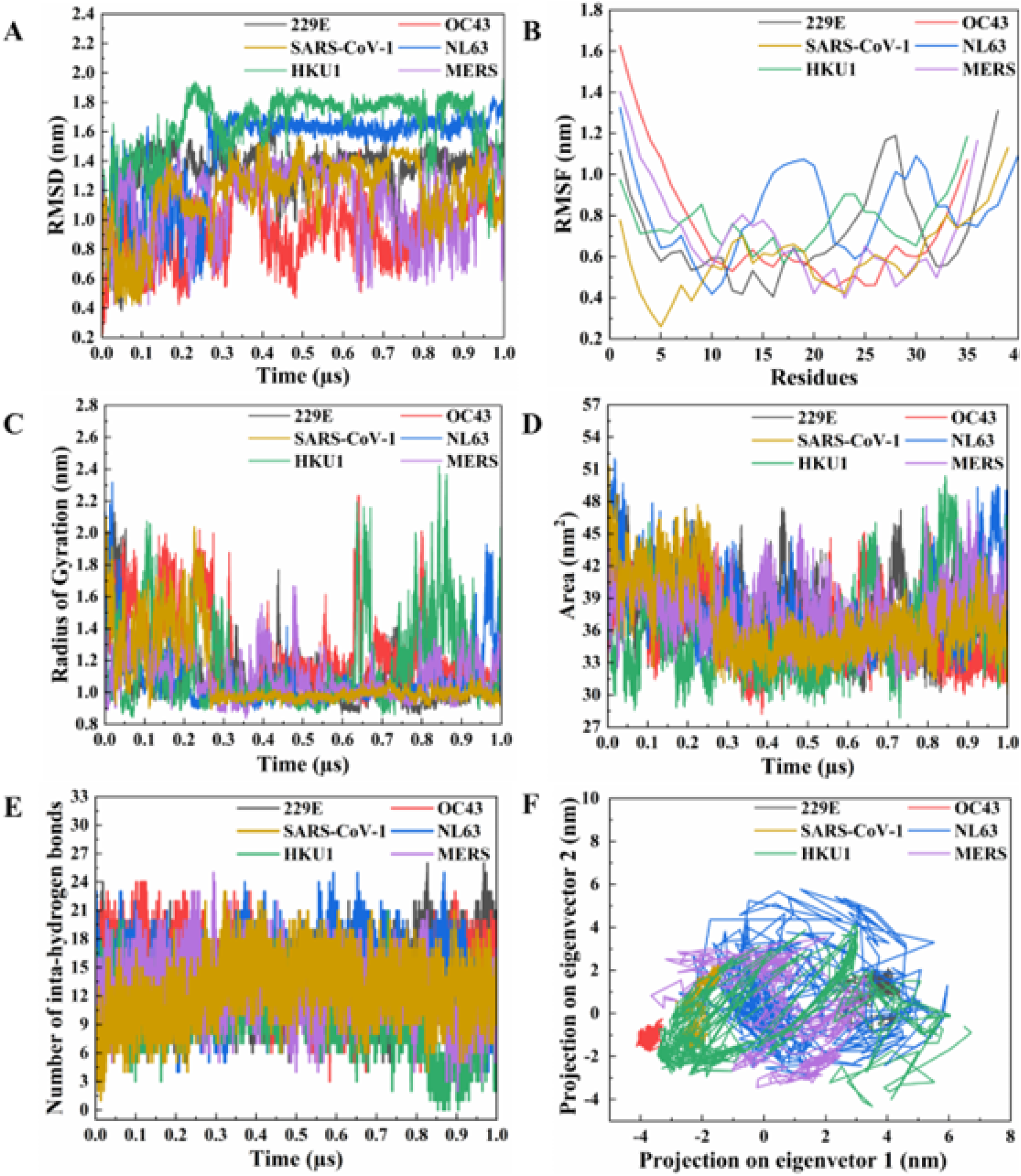
Molecular Dynamics Simulation trajectory investigation of the spike endodomains of all the six coronaviruses (229E, OC43, SARS-CoV-1, NL63, HKU1, MERS) for 1us, using CHARMM36m force field from Gromacs simulation package. (A) Root Mean Square Deviation (RMSD) (B) Root Mean Square Fluctuations (RMSF) (C) Radius of Gyration (RoG) (D) Principal Component Analysis (PCA) (E) Intra-hydrogen bonding (F) Solvent Accessibility Surface Area (SASA) plots over the simulation period.

### Analysis of Structural dynamics via MD simulations

From our previous study on the SARS-CoV-2 spike endodomain, we infer the structural disorderness of this region via modelling and simulation studies ^15^. Similarly, in our present study, we have checked the pattern of structural dynamics in spike endodomains of six other known coronaviruses. For this purpose, we have employed extensive simulations of 1 μs using AlphaFold2 modelled structure of spike endodomain. **Figure 7** shows the modelled structures of all six CoV endodomains. The rainbow-colored structures illustrate the structured and unstructured regions in the sequence of their chronological outbreaks. After preparing these structures for optimum symmetry and proper electrostatic distribution, the MD simulations are performed using CHARMM36m forcefield in the Gromacs package. Next, the solvation models are prepared using the CHARMM-GUI web server and processed further using our in-house HPC facility. Upon retrieving the solvated systems, the production run for 1 μs is done after minimization and equilibration.

As shown in **Figure 9**, there are significant structural transitions from ordered to disordered states in half and a total time of simulations. The structures at different frames with a periodic time difference of 50 ns are represented in **supplementary figures 2-7**. Also, these structures are superimposed on the modelled structures, which have shown tremendous RMSD values in SARS-CoV-1, NL63, HKU1, and MERS, illustrating their structural transitions during the simulation course. However, in the case of 229E and OC43, the structured region as a helix is majorly retained even after a 1 μs period.

**Figure 9:**
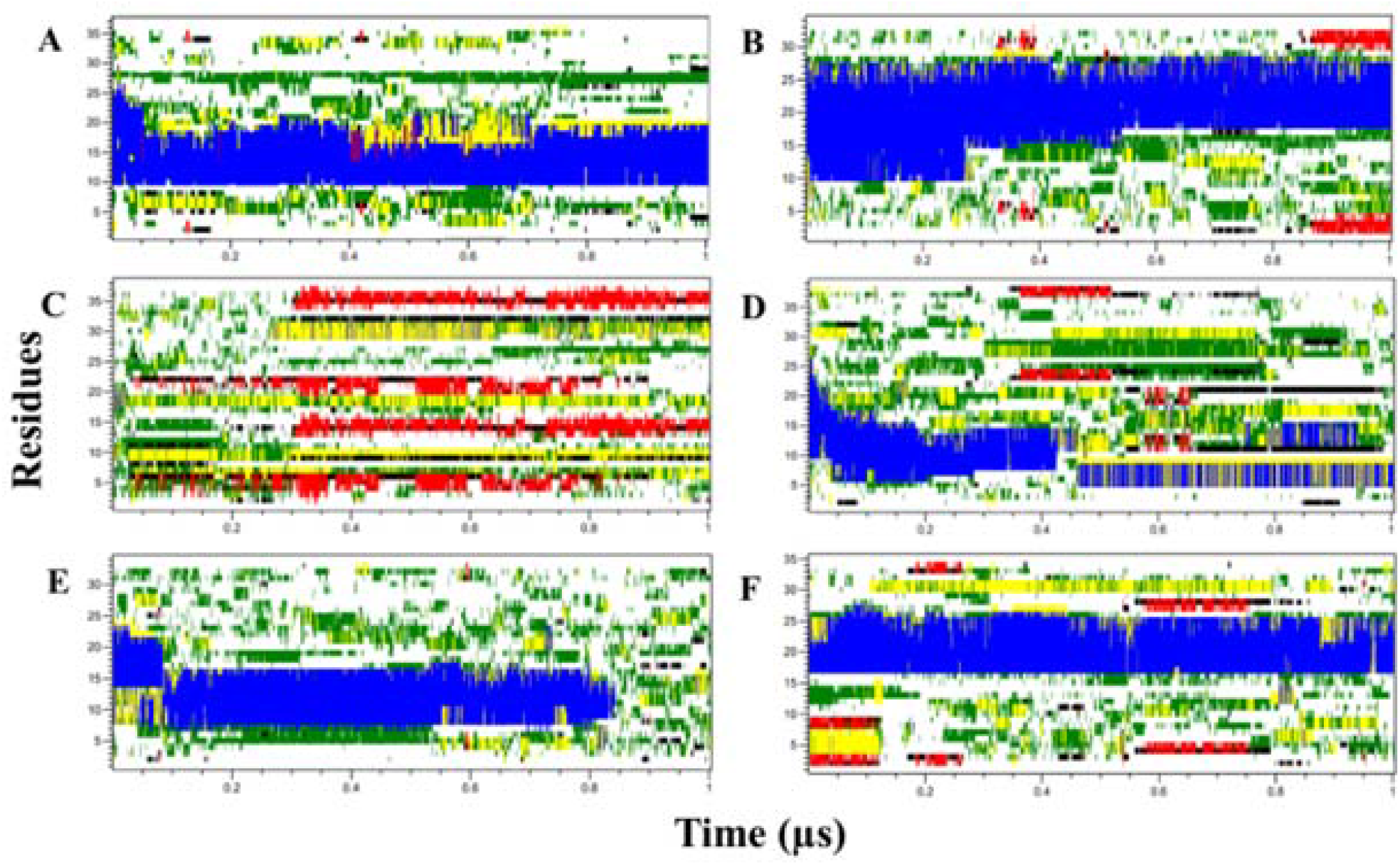
Timeline for the presence of secondary structures in the six coronaviruses, (A) 229E, (B) OC43, (C) SARS-CoV-1, (D) NL63, (E) HKU1, and (F) MERS, in accordance with simulation time of 1μs, using CHARMM36m forcefield from Gromacs simulation package. The color codes are as follows: beta sheets (red), beta bridge (black), bend (green), turn (yellow), alpha helix (blue), 3-helix (grey), and white blank area represent the coil region.

### 229E spike endodomain (residues 1136-1173)

No significant change was observed in structural composition during the simulation of 1 μs in 229E spike endodomain structure, as depicted by its structure in **Figure 7** and its trajectory analysis from **Figure 8**. The analyzed trajectory gives a maximum RMSD value of up to 1.75 nm with an average RMSD of ∼1.5 nm, and the residual fluctuations are shown by RMSF ranging between 0.3 nm to 1.3 nm. Moreover, the Rg revealed the compacted structure with the graph scale lying between 0.9 to 1.5 nm for the last 400 ns. It also forms an average of 12 hydrogen bonds within its structure for secondary structure stability. Further, we have used the initial two eigenvectors to estimate the overall dynamics of the protein in the provided system. Since these first two vectors are the main contributors to understanding the whole dynamics of the atomic coordinates of the protein, these vectors were plotted as eigenvetor1 on X-axis and eigenvector2 on Y-axis for the Principal component analysis (PCA). From the secondary structure timeline **(Figure 9)**, it was pretty clear that, more often, the simulated structure attains coil (white) and α-helical (blue) conformations. This was followed by bends (green) and turns (yellow). The ten residues at positions 10 – 20 were consistently showing α-helical conformation. Moreover, the rigidity of the structure can be assessed through the surface area of spike endodomain, which remains nearly similar to the initial with few fluctuations during simulations **(Figure 9D)**. The comparative secondary structure histogram depicted in Figure 10 showed more coils and 20% of α-helical conformations.

**Figure 10:**
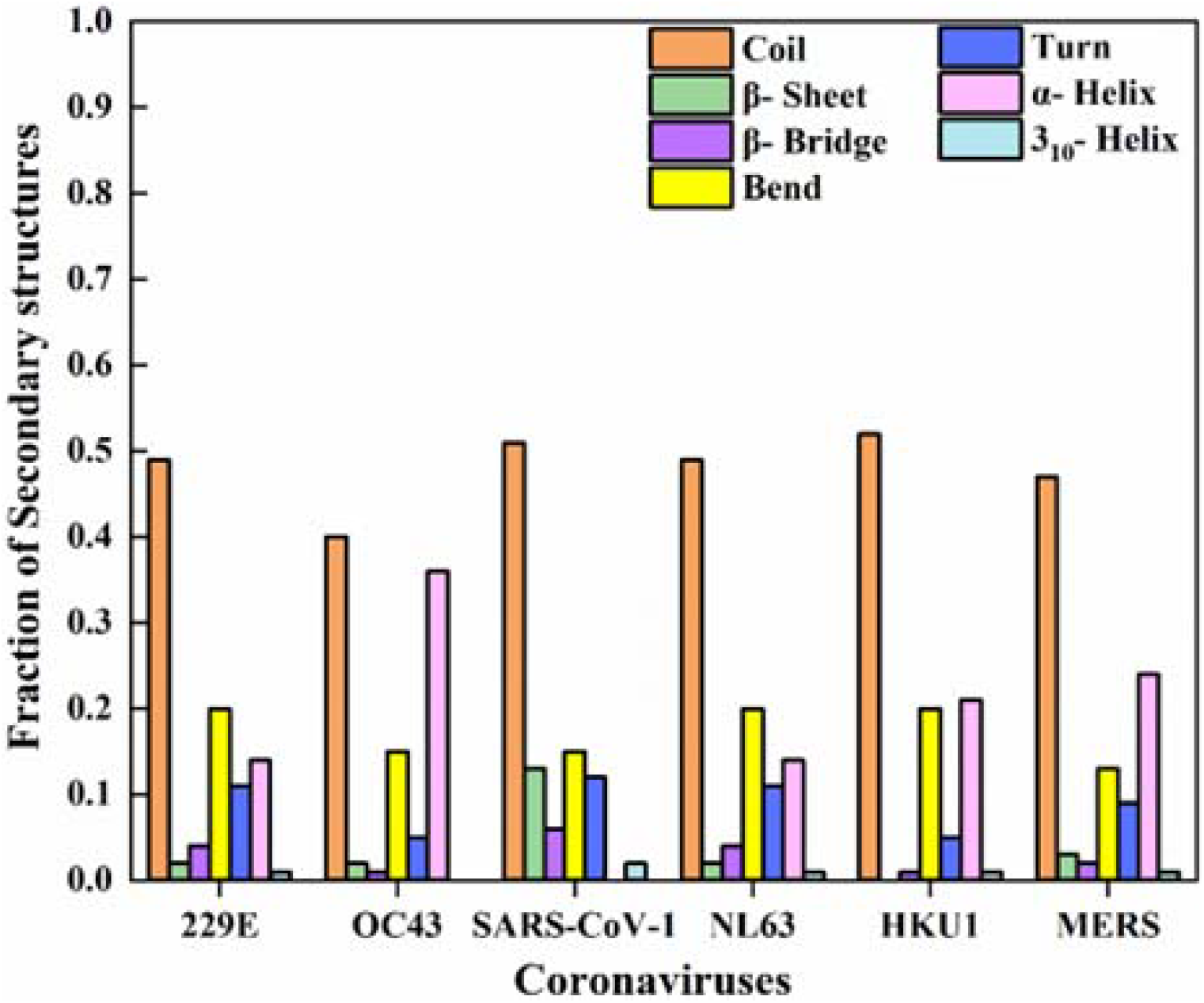
Identifying the extent of secondary structures present in the simulated models of the spike endodomains (229E, OC43, SARS-CoV-1, NL63, and HKU1) at 1us using the CHARMM36m force field from Gromacs simulation package.

### OC43 spike endodomain (residues 1319-1353)

However, in the case of OC43, 1325-1330 residues lose their helical conformations and are converted into the disordered form after one μs of the simulation run. The protein’s structural deviation for Cα from the reference starting structure gives the average value of 0.8 nm. But the RMS fluctuation lies between 0.5 nm to 1.6 nm for residues 1-10, and for the following residues viz 11-30, the values fall to near 0.5 nm while for the last ten residues, the value shoots up to 1.25 nm. This showed the disorderness propensity for both terminals of the endodomain residues. The gyration parameter fluctuates between 1.0 to 2.0 nm, which denotes less structural compactness to the completion of the simulation. The average number of 15 intra-hydrogen bonds maintained its helical structural conformation in some regions. Its structural conformation from the secondary structure timeline in **Figure 9** and histogram representation for the extent of the secondary structure **(Figure 10)** correlate well with the above results. From this, it is observed that nearly 40% fraction of the entire secondary structure forms α-helical conformation lying within residues 11-20 and is maintained throughout the simulations. The average surface area during the simulation run is ∼40 nm^2^, with some sharp fluctuations observed during half-time.

### SARS-CoV-1 spike endodomain (residues 1217-1255)

Unlike other modelled structures, the SARS-CoV-1 has an overall disordered structure, as observed from AlphaFold2 modelling. F**igure 7** gives a clear comparison of its modelled structure with its simulated structure as well as with other modelled structures. Its trajectory depicts a heavily fluctuating RMSD value approaching ∼1.6 nm for the entire trajectory. Here the residues fluctuate between the RMSF values of 0.2 and 1.3 nm. The radius of gyration also shows that the structure is quite fluctuating and remains disordered throughout the simulation course. There are, on average, 11 intra-hydrogen bonding that is formed during one μs simulation time. The other parameters, like Rg and SASA, have also shown fluctuations till 300 ns and then stabilized at a disordered conformation till the entire simulation time. **Figure 10** shows the structural conformations formed during simulation, showing that it has more coiled conformation (>50%) than the other structural elements.

### NL63 spike endodomain (residues 1317-1356)

The investigation for the dynamics of NL63 endodomain from Figure 7 shows that at the end of the simulation, the structure has loosened its integrity compared to the modelled helical structure. The trajectory also demonstrates its structural change at the end of the simulation as the RMSD for the last 300 ns lies between 1.5 nm to 2.0 nm. The fluctuation for individual residues varies from 0.5 nm to 1.25 nm, and the gyration also gives similar results with an average value of 1.5 nm for the last 400 ns with heavy fluctuations. The same trend has been observed in SASA, where the area increases and the number of intra-hydrogen bonding as the average number of bonding decreases as simulation time increases. As observed in the secondary structure timeline, a mixed population of all kinds of structures is seen and finally shows a disordered-like form of endodomain.

### HKU1 spike endodomain (residues 1322-1356)

Like NL63 endodomain, the HKU1 endodomain has also shown structural transitions after one μs simulation compared to its modelled structure where approx. 30% of the helical structure is present. As per the simulation results, the structure of NL63 endodomain started losing its structure halfway through the simulation. The helical propensity is lost during the last 200 ns and attains a disordered form. According to the 2struc server calculation, HKU1 has more than 50% of coiled regions and little portions of bend and helices. These outcomes are comprehended by the statistical trajectory analysis, where the mean deviation of atoms is highly varying and consists of values up to ∼2 nm. Similarly, the RMSF for all residues is in a heavily fluctuating region up to ∼1.2 nm. The change in hydrogen bonds and surface area is also noticeable where the variation in hydrogen bonds and the increase in SASA occurred.

### MERS spike CTR (residues 1318-1353)

The modelled structure of the endodomain of MERS has also shown a nearly 10% of secondary structure as calculated through the 2struc server. The initial structure consisted of a small helical region available throughout the simulation (**Figure 7**). As seen in the trajectory, the compactness of the MERS endodomain has remained nearly uncompromised as the Rg values fluctuate between 1.0 to 1.4 nm. Therefore, fluctuations are observed in only a few residues with no ordered structure. RMSD variations are felling between the range of 0.6-1.4 nm, which illustrates the deviation in the atoms in multiple frames during simulation. MERS endodomain has been observed to be present in the group of most ordered peptides having significant helical propensity like other members such as OC43. The PC analysis of the last 50 ns trajectory has validated the above results by showing comparatively less scattered plots than other peptides like SARS-CoV-1 and NL63.

### Free energy surface analysis

The free energy surface (FES) diagrams representing all populations present in the simulation trajectory are shown. First, the FES maps are plotted using two variables viz. RMSD and Rg. Then, using these variables, all frames are screened and mapped as per their calculated dG value. The presence of most dense regions depicts structures of similar energy. Here, we have shown three different structures from different map regions. According to the plots in **Figures 11A, 11B, and 11E**, most frames are clustered at one region, mainly ordered with lower Rg, while few structure frames with less ordered structures are dispersed at different regions with higher Rg. This shows that trajectory forms similar kinds of structures with the initial structure.

**Figure 11:**
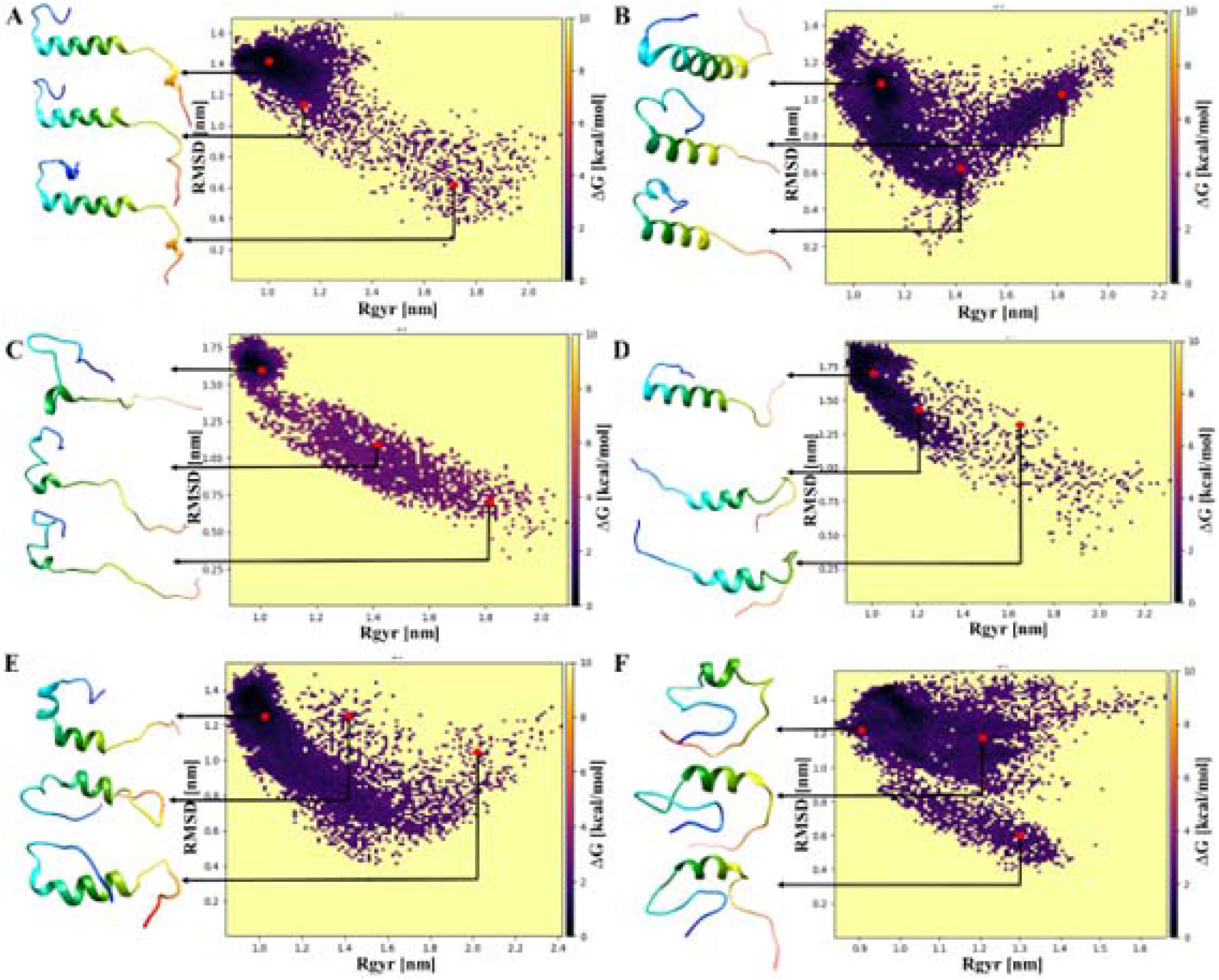
Free energy surface plots from the ordered parameters Rgyr [X-axis] and RMSD [Y-axis] depicting the different frames of the structures in order of their reduced free energies. (A) 229E, (B) OC43, (C) SARS-CoV-1, (D) NL63, (E) HKU1, and (F) MERS. The colour of the dots in the plot represents the ΔG [kcal/mol] values i.e., different feasible frames.

On the contrary, **11C and 11D** have scattered plots with varying populations at higher Rg depicting less ordered structures. Focusing on **11F**, i.e., MERS endodomain, it has shown two clustered populations at different RMSD and Rg. Finally, the third frame shown in **Figure 8** demonstrates that structures are nearly identical.

## Discussion

The Coronavirus Spike protein is the primary protein that encounters the host receptors. Generally, the structural changes for such interactions are crucial. The receptor binding domain of spike protein attains both open and closed forms during the interaction, where structural changes are observed ^15,34^. The glycosylated sites in spike glycoprotein 35 also regulate these structural changes. The spike’s structured domains, viz. extravirion domain, are highly studied and well explored in structure-function relationships. However, the transmembrane domain spanning around 20 residues and the following intravirion region is not explored for their structure-function or activity relationship. In our previous report on the SARS-CoV-2 spike cytoplasmic region, we deduced that this region is intrinsically disordered and contains gain of structure characteristics when exposed to some helix propensity enhancing solvents ^15^.

Not much information is available on disorder tendencies of the endodomain of Spike protein of the other six coronaviruses. Even though there is evidence for it to have such behavior as showcased by electron microscopy, where it is designated unmodelled due to variable conformation. Our previous publication has proven these assumptions on the SARS-CoV-2 virus via various bioinformatic and experimental approaches ^15^. An interesting phenomenon was observed after modelling the spike protein endodomains of the rest six coronaviruses using AlphaFold2: disordered propensity in the structure increased in the variants as they emerged. For instance, SARS-CoV-1, a highly virulent strain having approximately 70% of its residues disordered, whereas strains like 229E or NL63, which are comparatively less virulent as per records, have disorders of approximately 15 and 30%, respectively.

Further, the modelled 3D structure analysis also suggests the same. The snapshots of the one μs MD simulation trajectory have revealed that no significant conformational structure change occurred during simulation for less virulent strains like 229E or OC43. However, the amount of conformational change is much more significant in the case of more virulent strains like SARS-CoV-1 or MERS. Moreover, the highly virulent variant SARS-CoV-2, the endodomain, was completely disordered after simulation ^15^. Moreover, the full-length spike proteins modelled structures have shown a very low propensity for the helical or beta sheets in intravirion regions or endodomains which majorly constitutes random coil in their structures.

Various studies previously performed on disordered proteins have widely demonstrated their ability to bind with multiple downstream substrates ^36,37^. This trait of promiscuous binding can be critical for organisms with small genomes like viruses, allowing a single protein to perform multiple functions ^38^. There are a few reports linking disorder in the viral genome and how it affects host pathogenicity ^39^, but no such studies have been done linking disorder with the specific regions of viruses and pathogenicity. However, the cytoplasmic tails at C-terminal regions have been observed to be involved in viral assembly and pathogenicity. Like in the case of Measles, the cytoplasmic tails of glycoproteins hemagglutinin (H) and fusion protein (F) are involved in viral assembly and cell-to-cell fusion, and their deletion caused variable effects on receptor binding and cell fusion promotion activity ^40^. This study aligns with a similar study on SARS-CoV, where changes in the composition of the C-terminal residues have correlated with changes in its pathogenic properties ^13^. In another case of Influenza, type A and B viruses possess Hemagglutinin (HA) and Neuraminidase on their surface. The virus infects the host cell in a complex multistep process involving cleavage of the Hemagglutinin by cell proteases to reveal a fusion peptide that inserts itself into the endosome. At the same time, the extreme C-terminal region helps anchor the viral membrane during the process ^41^. Hence, these extreme C-terminal regions of viral proteins may be necessary for their pathogenesis and infectivity. The functional relevance of the endodomain of coronaviruses has not been well studied. As observed in our data, correlating the disorder tendency of the S protein endodomain and viral infectivity might provide an important area to explore in the future.

A few subcellular localization motifs are also present on the endodomain of spike, i.e., COPI binding KXHXX-type dibasic motif or the diacidic COPII binding DEDDSE motif. Our search for other eukaryotic motifs on spike endodomain led to an exciting revelation of the presence of multiple phosphorylation motifs on the endodomain or C-terminal tail of all coronaviruses, coinciding with palmitoylation regions as well ^17^. Palmitoylation of C-terminal tail cysteine residues has been previously studied to be implicated with viral pathogenicity ^13,29,42^. Furthermore, reversible phosphorylation and palmitoylation have been observed in certain eukaryotic proteins where it regulates transport. For example, in the case of protein phosphodiesterase 10A, phosphorylation at Thr-16 prevents palmitoylation at Cys-11, interfering with the subcellular localization of the protein ^30^. This phenomenon may also influence the post-translational modifications on spike C-terminal tail. According to the results from ELM server, we found multiple phosphorylation motifs in proximity to the palmitoylation motifs. We, therefore, can propose the same set of events to regulate the budding of S protein into the intermediate compartments between ER and Golgi. Also, it may affect its ability to interact with the M protein, which is another structural protein necessary for viral assembly, and thereby influence viral propagation.

## Conclusion

After the outbreak in December 2019 in Wuhan, China, many studies have been published on other CoVs, considering the need to understand the newly emerged SARS-CoV-2. Since spike glycoprotein is the first and foremost protein to interact with the host receptor upon infection, it is imperative to dig out its structure, functions, and related properties. However, none of the available structures of spike glycoprotein of CoVs has the transmembrane and the extreme C-terminal region or endodomain. Therefore, without the three-dimensional structure of spike glycoprotein endodomain, we have explored these regions through multiple bioinformatics approaches. Using sequence and structure-based analysis of these regions, we have identified several molecular recognition features (MoRFs), which lie in the disordered regions, the short linear motifs, and the possible post-translational modification sites in those regions. Also, the structural conformations for their extreme dynamics have been studied through MD simulations. These outcomes of all six CoVs and previous revelation on SARS-CoV-2 spike-endodomain may help to understand the holistic perspective of intraviral protein interactions of spike and the host proteins during infection.

## Material and methods

### Sequence retrieval and Multiple sequence alignment

Spike endodomain sequences were retrieved from full-length spike S2 subunit sequences from UniProt, annotated under the High-quality Automated and Manual Annotation of Proteins (HAMAP) rule. Multiple sequence alignment (MSA) of these sequences was done using the ClustalΩ program, which is fast, scalable, and generates high-quality sequence alignment. It uses a modified mBed method to produce accurate guide trees aligned via Hidden Markov Model (HMM) to generate accurate alignments ^43^. ESPript 3.0 (Easy Sequencing in PostScript) was used to create aligned sequence images ^44^.

### Disorder Prediction

To reduce the prediction biasedness, various webservers were employed to predict the tendency of the cytoplasmic tail of spike proteins to disorder. Disorder predictors from Predictor of Natural Disordered Regions (PONDR) family, i.e., PONDR^®^ FIT ^45^, PONDR^®^ VLXT ^46^, PONDR^®^ VL3 ^47^, PONDR^®^ VSL2^48^, along with IUPred2A^49^, PrDOS ^50^ and DisEMBL ^51^ were used. In addition, the redox function of IUPRED2A was employed due to the cysteine-rich cytoplasmic domain. Residues with a score >0.5 were considered disordered residues. The percent of predicted intrinsic disorder (PPID) from the output of all predictors for cytoplasmic regions of all the spike proteins was calculated as follows:

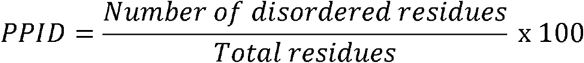

### Molecular Recognition Feature (MoRF) and Short Linear Motifs (SLiMs) Prediction

Molecular recognition features (MoRFs) are small binding regions within proteins around (5-25) residues in length, located inside disordered regions^52^. Our study used three servers to predict MoRFs in spike endodomains, including MorfCHiBi_Web^53^, Anchor2^49^, and Disopred 3^54^. For the MorfChiBi_Web server, the threshold value was kept at 0.725, for Anchor2, Disopred 3, the threshold value was kept at 0.5.

SLiMs, also known as short linear motifs or MiniMotifs, have 3-11 residues and are the functional regions present in intrinsically disordered regions, and the interactions mediated by these have been reported to regulate various processes. Therefore, the cytoplasmic spike region of all coronaviruses was searched for functional annotation using the eukaryotic linear motif (ELM) server 55, which provides a manually curated data set with experimentally validated SLiM instances ^56^.

### Post-translational modification prediction

We have also predicted the residues within the endodomains that are prone to post-translational modifications using MusiteDeep server^57^. The server predicts a total of 9 types of modifications, viz. Phosphorylation, Glycosylation, Ubiquitination, Sumoylation, Acetylation, Methylation, Pyrrolidone carboxylic acid, Palmitoylation, and Hydroxylation.

### Structure Modelling

Structure models of all six spike full-length proteins and their endodomains of coronaviruses were modelled using AlphaFold2^58^. It is a highly accurate and advanced tool that uses an artificial intelligence-based approach and generates 3D protein structures with near experimental accuracy. For secondary structure component percentage calculation, the *2Struc* server was used^59^.

### Molecular dynamics simulations

Molecular dynamics simulations were performed to determine the conformational changes in the modelled structure of the spike endodomain of all six coronaviruses. Here, the solvated input files for simulations were generated using CHARMM-GUI v3.5 (Chemistry at HARvard Macromolecular Mechanic-Graphical User Interface)^60^. It uses CHARMM36m potential energy function (forcefield) program to prepare the system’s atomic coordinates and analyze the trajectory. The solution builder module produced the final input files by providing the aqueous solvent environment to the provided protein structure. TIP3P solvent model solvated the system in a rectangular water box.

Moreover, the protein was centered at a 10Å edge distance from the box. After this, the whole system was neutralized with the Monte-Carlo ions placement method. Particle mesh Ewald-fast Fourier transform (PME-FFT) was used for the periodic boundary condition’s grid generation. The ensembles NVT for the equilibration input and NPT for the dynamics input was set with the temperature of 303.15 K. All the input files were downloaded for the Gromacs package^61^. The energy minimization of the system was done by running the steepest descent method for 50,000 steps. Then the system was equilibrated using Nose-Hoover temperature coupling and Parrinello-Rahman pressure coupling methods for 125 ps. Finally, MD production was run for one μs using the in-house HPC facility using Gromacs v5^62^. The Verlet cut-off scheme and LINCS constraint algorithm were used for the energy minimization, equilibration, and production steps. Different parameters were taken for the trajectory analysis, i.e., Root Mean Square Deviation (RMSD), Root Mean Square Fluctuation (RMSF), Radius of Gyration (RoG), intra-hydrogen bonding, Principal Component Analysis (PCA), Free Energy Surfaces (FES), and Solvent accessible surface area (SASA).

## Supplementary Material Description

The supplementary file is attached. It contains the disorder profiles and SLiM information of all CoVs spike endodomains.

## Acknowledgments

All the authors would like to thank the HPC facility, IIT Mandi, for the infrastructure. RG is thankful to the Government of India for the grants, including IYBA award (BT/11/IYBA/2018/06), MHRD-SPARC (SPARC/2018-2019/P37/SL), and Science and Engineering Research Board (SERB), India (Grant Number: CRG/2019/005603), and Indian Council of Medical Research (58/6/2020/PHA/BMS, and 52/04/2020/BIO/BMS). AB, BM, and RJ are thankful for their JRF from MHRD, Govt. of India.

## Author contribution

RG: Conception, design & study supervision. PK, AB, BM, and RJ: acquisition and interpretation of data. PK, AB, BM, and RJ: contributed to paper writing.

