## Supplementary Figure and Supplementary Table for "Coronaviruses Spike glycoprotein endodomains: the sequence and structure-based comprehensive study"

**Supplementary Information**

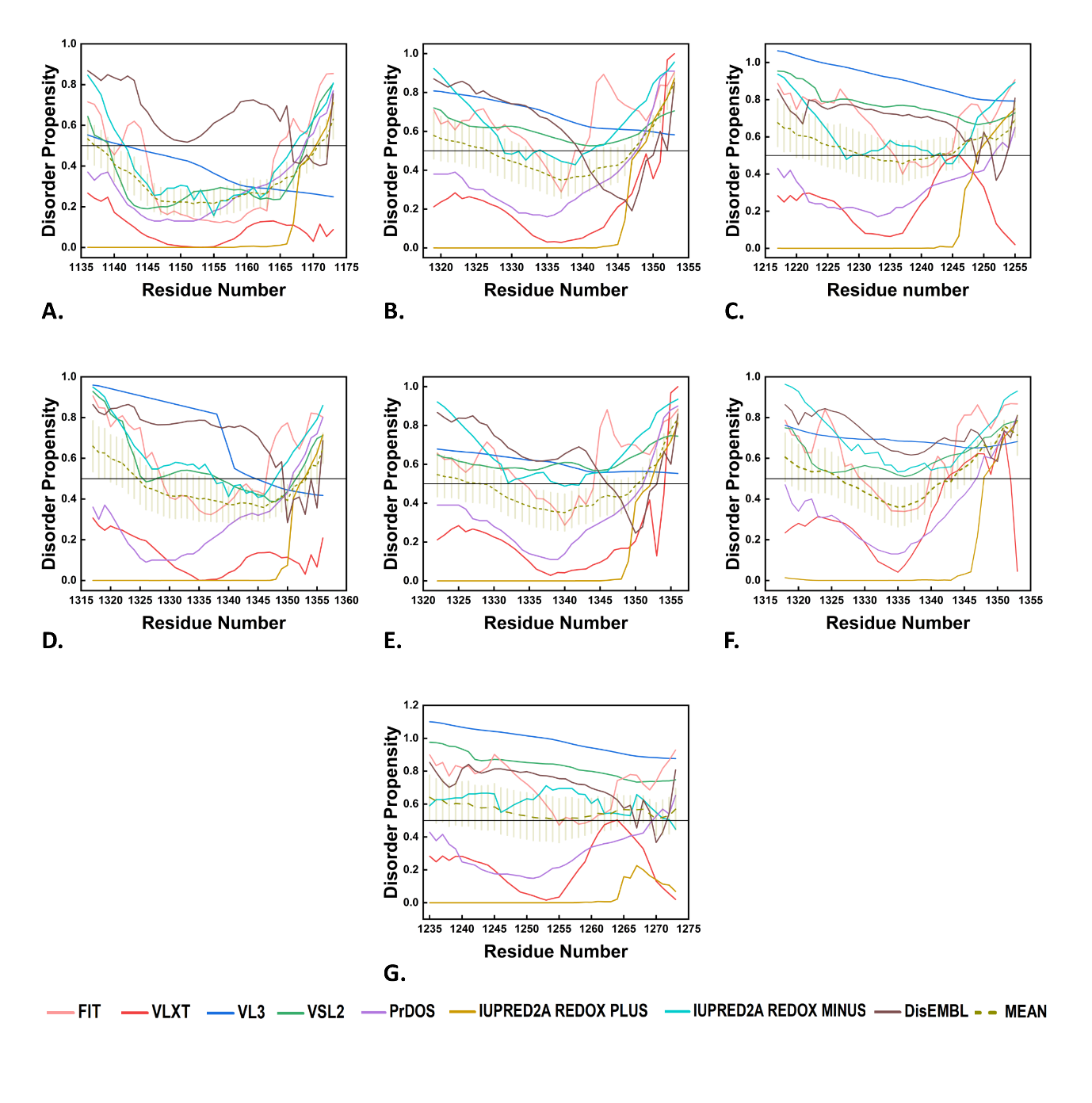

**Supplementary Figure 1:** Intrinsic disorder analysis of spike C-terminal cytoplasmic tail of 6 human coronaviruses: from top left A. 229E, B. OC43, C. SARS- CoV-1, D. NL63, E. HKU1, F. MERS and G. SARS-CoV-2 using seven predictors including PONDR family (FIT, VL3, VLXT, VSL2), IUPred 2A (Redox plus and Redox minus), DisEMBL, and PrDOS servers. For the comparison purpose, we are also showing our previously studied, SARS-CoV-2 spike CTR prediction using the similar servers ^1^.

**Supplementary Table 1**: Identified Short Linear Motifs (SLiMs; 3-10 long amino acid residues) in human coronaviruses spike endodomains: MERS, HKU1, OC43, NL63, 229E, SARS-CoV-1, and SARS-CoV-2:

| **Name** | **Motif Sequence** | **Motif Sequence Number** | **Identified SLiMs** | **SLiMs Descriptions** |
| --- | --- | --- | --- | --- |
| **MERS** | DRYEEY | 1338-1343 | LIG_LIR_Nem_3 | Nematode-specific variant of the canonical LIR motif that mediate processes involved in autophagy. |
|  | YDLEP | 1343-1347 | LIG_TRFH_1 | The TRFH binding motifs are found in proteins recruited to sheliterin, which protects the mammalian telomers, by TRF1 and TRF2 |
|  | LCCTGCGT | 1318-1326 | MOD_GSK3_1 | GSK3 phosphorylation recognition site |
|  | NRC | 1335-1337 | MOD_N-GLC_2 | Atipical motif for N-glycosylation site. |
|  | LCCTGC | 1318-1323 | MOD_NEK2_1 | NEK2 phosphorylation motif |
| **HKU1** | GSACF | 1328-1332 | LIG_BRCT_BRCA1_1 | Phosphopeptide motif which directly interacts with the BRCT (carboxy-terminal) domain of the Breast Cancer Gene BRCA1 with low affinity |
|  | GSACFSK | 1328-1334 | LIG_BRCT_BRCA1_2 |  |
|  | IKTSHDD | 1350-1356 | LIG_FHA_2 | Phosphothreonine motif binding a subset of FHA domains |
|  | GSAC | 1328-1331 | MOD_GlcNHglycan | Glycosaminoglycan attachment site |
|  | CCCTGCGS; GCGSACFS | 1322-1329; 1326-1333 | MOD_GSK3_1 | GSK3 phosphorylation recognition site |
|  | NCC | 1337-1339 | MOD_N-GLC_2 | Atipical motif for N-glycosylation site. |
| **OC43** | IKTSHDD | 1347-1353 | LIG_FHA_2 | Phosphothreonine motif binding a subset of FHA domains |
|  | TGYQELVI | 1340-1347 | LIG_LIR_Gen_1 | Canonical LIR motif that binds to Atg8/LC3 protein family members to mediate processes involved in autophagy. |
|  | DDYTGY  DYTGY  TGYQEL | 1337-1342; 1338-1342; 1340-1345 | LIG_LIR_Nem_3 | Nematode-specific variant of the canonical LIR motif that mediate processes involved in autophagy. |
|  | YTGY | 1339-1342 | LIG_SH2_STAT5 | STAT5 Src Homology 2 (SH2) domain binding motif. |
|  | CCCTGCGT | 1319-1326 | MOD_GSK3_1 | GSK3 phosphorylation recognition site |
|  | CGTSCF | 1324-1329 | MOD_NEK2_2 | NEK2 phosphorylation motif |
|  | CTGCGTSC | 1321-1328 | MOD_OFUCOSY | Site for attachment of a fucose residue to a serine. |
|  | YQEL | 1342-1345 | TRG_ENDOCYTIC_2 | Tyrosine-based sorting signal responsible for the interaction with mu subunit of AP (Adaptor Protein) complex |
| **229E** | FFSCF | 1145-1149 | LIG_Pex14_2 | Fxxx[WF] motifs are present in Pex19 and S. cerevisiae Pex5 cytosolic receptors that bind to peroxisomal membrane docking member, Pex14 |
|  | LPYYDV | 1162-1167 | LIG_TYR_ITIM | ITIM (immunoreceptor tyrosine-based inhibitory motif). Phosphorylation of the ITIM motif, found in the cytoplasmic tail of some inhibitory receptors (KIRs) that bind MHC Class I, leads to the recruitment and activation of a protein tyrosine phosphatase. |
|  | GFFSCFAS | 1144-1151 | MOD_GSK3_1 | GSK3 phosphorylation recognition site |
|  | FASSIR | 1149-1154 | MOD_NEK2_1 | NEK2 phosphorylation motif |
|  | GFFSCF | 1144-1149 | MOD_NEK2_2 | NEK2 phosphorylation motif |
|  | CCGFFSC  CGFFSC | 1142-1148  1143-1148 | MOD_OFUCOSY | Site for attachment of a fucose residue to a serine. |
|  | YYDV | 1164-1167 | TRG_ENDOCYTIC_2 | Tyrosine-based sorting signal responsible for the interaction with mu subunit of AP (Adaptor Protein) complex |
| **NL63** | EFEKVHV | 1349-1355 | LIG_LIR_Gen_1 | Canonical LIR motif that binds to Atg8/LC3 protein family members to mediate processes involved in autophagy. |
|  | EFEKV | 1349-1353 | LIG_LIR_Nem_3 | Nematode-specific variant of the canonical LIR motif that mediate processes involved in autophagy |
|  | LTSSMR | 1330-1335 | MOD_NEK2_1 | NEK2 phosphorylation motif with preferred Phe, Leu or Met in the -3 position to compensate for less favorable residues in the +1 and +2 position. |
|  | YYEF | 1347-1350 | TRG_ENDOCYTIC_2 | Tyrosine-based sorting signal responsible for the interaction with mu subunit of AP (Adaptor Protein) complex |
| **SARS-CoV-1** | KGVKL | 1348-1352 | CLV_PCSK_SKI1_1 | Subtilisin/kexin isozyme-1 (SKI1) cleavage site ([RK]-X-[hydrophobic]-[LTKF]-\|-X). |
|  | SCCSCLK  SCGSCCK | 1221-1227  1331-1337 | MOD_CK1_1 | CK1 phosphorylation site |
|  | CCMTSCCS | 1217-1224 | MOD_GSK3_1 | GSK3 phosphorylation recognition site |
|  | CCMTSC  CMTSCCSC  CLKGACSC  CSCGSC | 1217-1222  1218-1225  1225-1332  1230-1235 | MOD_OFUCOSY | Site for attachment of a fucose residue to a serine. |
| **SARS CoV-2** | KGVKL | 1266-1270 | CLV_PCSK_SKI1_1 | Subtilisin/kexin isozyme-1 (SKI1) cleavage site ([RK]-X-[hydrophobic]-[LTKF]-\|-X). |
|  | SCCSCLK  SCGSCCK | 1239-1245  1249-1255 | MOD_CK1_1 | CK1 phosphorylation site |
|  | CCMTSCCS | 1235-1242 | MOD_GSK3_1 | GSK3 phosphorylation recognition site |
|  | CCMTSC  CMTSCCSC  CLKGCCSC  CCSCGSC  CSCGSC | 1235-1240  1236-1243  1243-1250  1247-1253  1248-1253 | MOD_OFUCOSY | Site for attachment of a fucose residue to a serine. |
